# Non-random mating between nesting sites of Hawaiian hawksbill turtles: demographic discontinuity within a small isolated population

**DOI:** 10.1101/2022.10.30.514389

**Authors:** John B. Horne, Amy Frey, Alexander R. Gaos, Summer Martin, Peter H. Dutton

**Affiliations:** Southwest Fisheries Science Center, NOAA-Fisheries, La Jolla, California, USA; This research was performed while the author held an NRC Research Associateship Award from the National Academies of Science, Engineering, and Medicine; Pacific Islands Fisheries Science Center, NOAA-Fisheries, Honolulu, Hawaii, USA

**Keywords:** conservation genetics, sea turtles, breeding sex ratios, nesting frequency, inbreeding

## Abstract

Hawksbill sea turtles (*Eretmochelys imbricata*) from the Hawaiian archipelago form a small, genetically isolated, population consisting of only a few tens of individuals breeding annually. Most females nest on the island of Hawai’i, but little is known about the demographics of this rookery. This study used genetic relatedness, inferred from 135 microhaplotype markers, to determine breeding sex-ratios, estimate female nesting frequency, and assess relationships between individuals nesting on different beaches. Samples were collected during the 2017 nesting season and final data included 13 nesting females and 1,002 unhatched embryos, salvaged from 41 nests, 13 of which had no observed mother. Results show that most females used a single nesting beach laying 1-5 nests each. From female and offspring alleles the paternal genotypes of 12 breeding males were reconstructed and many showed high relatedness to their mates. Pairwise relatedness of offspring revealed one instance of polygyny but otherwise suggest a 1:1 breeding-sex ratio. Relatedness analysis and spatial-autocorrelation of genotypes indicate non-random mating among complexes of nesting beaches, for both sexes, suggesting strong natal philopatry. Nesting complexes also showed unique patterns of inbreeding and outbreeding across loci, further indicating that Hawaiian hawksbill turtles have demographically discontinuous nesting populations at a fine spatial scale.

## Introduction

It is increasingly recognized that the maintenance of a population’s size and corresponding genetic diversity is necessary for its long-term persistence (Luque et al. 2016; Hoban et al. 2020; Kardos et al. 2021). Population decline often results in a significant loss of genetic diversity that leads to a cascade of detrimental effects such as inbreeding depression (Keller and Waller 2002; Reed et al. 2002; O’Grady et al. 2006; Jamieson and Allendorf 2012; Caballero et al. 2017; Hoban et al. 2020), loss of adaptive potential (Pinsky and Palumbi 2014; Hoffman et al. 2017), and a reduced capacity for population recovery (Hughes and Stachowicz 2004; Walsh et al. 2006; Worm et al. 2006; Johnson et al. 2016). For populations that remain small and isolated for prolonged periods of time the risk of extinction is high (Kardos et al. 2021).

Most sea turtle populations worldwide have been dramatically reduced in size from historical levels, and virtually all still face challenges to persistence posed by a variety of human-related impacts (Hamann et al. 2010; Wallace et al. 2011; Riskas et al. 2016; Jensen et al. 2018). The alarming rate of sea turtle decline has prompted several decades of conservation effort that has resulted in some cases of meaningful population recovery (Chaloupka et al. 2008; Hamilton et al. 2015; Valdivia et al. 2019; Pritchard et al. 2022), suggesting that proper management can reverse negative trends and help ensure long-term persistence. However, not all populations have responded well to conservation protections and it is currently unclear what role genetic diversity may play in the ability of some sea turtle populations to rebound after experiencing demographic bottlenecks (Komoroske et al. 2017).

The hawksbill turtle (*Eretmochelys imbricata*) is a circumtropically distributed marine reptile with nesting colonies scattered across the Atlantic and Indo-Pacific. In the past their numbers have dwindled for many of the same reasons as other sea turtle species but also for being disproportionately targeted by the international shell trade, which is illegally ongoing in many countries (LaCasella et al. 2021). Some hawksbill populations have shown signs of recent recovery (Hamilton et al. 2015; Valdivia et al. 2019) in response to management interventions, but others continue to decline despite efforts to protect their remaining numbers (Bell et al. 2020). Currently, *E. imbricata* is listed as endangered under the US Endangered Species Act (ESA; NMFS and USFWS 2013), and as critically endangered by the IUCN Red List (Mortimer and Donnelly 2008).

In the Hawaiian archipelago, hawksbill turtles are described as rare and their resident population is thought to include less than 100 adult females, with only 5-26 nesting annually (van Houtan et al. 2012; van Houtan et al. 2016; Gaos et al. 2021). The population may also be predominantly female, as only 1 in 5 strandings in Hawaii are male (Brunson et al. 2022). Field surveys of nesting activity have been conducted since the 1980s with varying levels of effort, and predictive modeling based on these data suggests a recent uptick in the number of nesting females (Gaos et al. 2021). Nevertheless, the population remains extremely small, even though hawksbills have had endangered species status in the United States since 1970 (https://www.federalregister.gov/citation/35-FR-8491).

Given the vulnerability of Hawaiian hawksbill turtles, due to their chronically low abundance, and the fact that they harbor unique mitochondrial genetic diversity for this species (Gaos et al. 2020), there is motivation to better understand the conservation needs of this population. Monitoring these turtles is difficult, however, because in addition to being one of the smallest sea turtle populations in the world (van Houtan et al. 2016) nesting also takes place at multiple remote and hard-to-access beaches, most of which are located along the southern coast of Hawai’i Island (Gaos et al. 2021). Consequently, many nests are discovered with unknown mothers and field efforts have been unable to confidently census nesting females. A consequence of these uncertainties is that the mating system of Hawaiian hawksbills, and their operational breeding sex ratios are unknown.

One way to fill demographic information gaps for Hawaiian hawksbill turtles is with genetic relatedness analysis. Using genetic markers, it is possible to infer parent-offspring relationships between known nesting females and nests where the mother is unidentified (Frey et al. 2013; 2014) and obtain a more complete population census. The genetic relationships of offspring within and among nests also indicate patterns of paternity, and can be used to obtain information on mating systems and breeding sex ratios (Stewart and Dutton 2011; 2014a; Lasala et al. 2018). In this study, an amplicon-based array of single nucleotide polymorphisms (SNPs), and small nucleotide insertion-deletion (indel) markers, was developed to capture population-specific variation in Hawaiian hawksbill turtles and to estimate pairwise genetic relatedness. The data presented hereafter provide insight into the basic reproductive biology of a precariously endangered population of sea turtles, and also show that such studies can yield unexpected results with importance to conservation management.

## Methods

### Sample collection and study design

Tissue samples from nesting females were collected during the 2017 breeding season at five nesting beaches located along the southern coast of Hawai’i island. Samples consisted of a skin biopsy (∼ 0.5 cm^2^) taken from the neck or shoulder. Tissue was stored in a high-salt solution for transport. Offspring tissue collections were flippers of dead embryos salvaged from 41 nests (Table 1) after live hatchlings had vacated the nesting chamber. For thirteen of these nests the mother was not observed during nesting and is unknown. Ten females from the 2018 nesting season were included in this study to act as control samples for assigning mothers to nests. Female Hawaiian hawksbills are not known to nest in consecutive years (Gaos et al. 2021), so it is presumed that these off-year individuals could not have been the unidentified mothers and thus serve as negative controls. Inconel flipper tags (National Band & Tag, Newport, KY, USA) and Passive Integrated Transponder (PIT; Avid, Norco, CA, USA) tags were applied to all female turtles encountered to confirm and track identity during the nesting seasons.

**Table 1:** Hawaiian hawksbill turtle nests from 2017, with the observed mother, inferred mother from parentage analysis, probability of maternity, the ID of the paternal genotype constructed from offspring genotypes, and the estimated relatedness coefficient between the mating pair (Lynch and Ritland 1999). Inferred mothers marked with an asterisk were not fully supported by all analyses. Note: no genetics sample of mother-160 was available for this study.

| Nest | Nesting complex | Number of hatchlings | Observed Mother | Inferred Mother | Prob. | Paternal Genotype ID | Mating pair r |
| --- | --- | --- | --- | --- | --- | --- | --- |
| Apua-01 | ‘Āpua | 26 | 153 | 153 | 1.000 | P01 | 0.25 |
| Apua-02 | ‘Āpua | 46 | 153 | 153 | 1.000 | P01 | 0.25 |
| Apua-03 | ‘Āpua | 4 | 155 | 155 | 1.000 | P02 | 0.14 |
| Apua-06 | ‘Āpua | 27 | 153 | 153 | 1.000 | P01 | 0.25 |
| Apua-07 | ‘Āpua | 33 | none | 153 | 1.000 | P01 | 0.25 |
| Apua-08 | ‘Āpua | 80 | 158 | 158 | 1.000 | P03 | 0.24 |
| Apua-09 | ‘Āpua | 55 | 154 | 154 | 1.000 | P04 | 0.15 |
| Apua-10 | ‘Āpua | 12 | 155 | 155 | 1.000 | P02 | 0.14 |
| Apua-11 | ‘Āpua | 30 | 153 | 153 | 1.000 | P01 | 0.25 |
| Apua-12 | ‘Āpua | 37 | none | 158 | 1.000 | P03 | 0.24 |
| Apua-14 | ‘Āpua | 42 | 155 | 155 | 1.000 | P02 | 0.14 |
| Apua-15 | ‘Āpua | 98 | none | 158 | 1.000 | P03 | 0.24 |
| Apua-16 | ‘Āpua | 23 | none | 158 | 1.000 | P03 | 0.24 |
| Halape-01 | ‘Āpua | 79 | none | 85 | 1.000 | P06 | 0.28 |
| Halape-02 | ‘Āpua | 10 | 85 | 85 | 1.000 | P06 | 0.28 |
| Halape-03 | ‘Āpua | 79 | 85 | 85 | 1.000 | P06 | 0.28 |
| Pohue-01 | Pōhue | 4 | none | 119 | 0.991 | P07 | 0.27 |
| Pohue-02 | Pōhue | 4 | 151 | 151 | 1.000 | P09 | -0.06 |
| Pohue-03 | Pōhue | 31 | 152 | 152 | 1.000 | P10 | -0.05 |
| Pohue-04 | Pōhue | 6 | 119 | 119 | 0.991 | P07 | 0.27 |
| Pohue-05 | Pōhue | 13 | 71 | 71 | 1.000 | P11 | 0.0 |
| Pohue-06 | Pōhue | 37 | 76 | 76 | 1.000 | P07 | 0.07 |
| Pohue-07 | Pōhue | 17 | 151 | 151 | 1.000 | P09 | -0.06 |
| Pohue-08 | Pōhue | 17 | 152 | 152 | 1.000 | P10 | -0.05 |
| Pohue-09 | Pōhue | 8 | none | 157 | 1.000 | P05 | 0.09 |
| Pohue-10 | Pōhue | 21 | 119 | 119 | 0.991 | P07 | 0.27 |
| Pohue-11 | Pōhue | 51 | 76 | 76 | 1.000 | P07 | 0.07 |
| Pohue-12 | Pōhue | 17 | 71 | 71 | 1.000 | P11 | 0.0 |
| Pohue-13 | Pōhue | 1 | 151 | 151 | 1.000 | P09 | -0.06 |
| Pohue-15 | Pōhue | 22 | 152 | 152 | 1.000 | P10 | -0.05 |
| Pohue-17 | Pōhue | 19 | 71 | 71 | 1.000 | P11 | 0.0 |
| Pohue-18 | Pōhue | 11 | 151 | 151 | 1.000 | P09 | -0.06 |
| Pohue-19 | Pōhue | 26 | 76 | 76 | 1.000 | P07 | 0.07 |
| Pohue-20 | Pōhue | 24 | none | 76 | 1.000 | P07 | 0.07 |
| Pohue-21 | Pōhue | 13 | 152 | 152 | 1.000 | P10 | -0.05 |
| Pohue-24 | Pōhue | 12 | none | 159* | 0.687 | P12 | -0.04 |
| Pohue-25 | Pōhue | 6 | 160 | 159* | 0.687 | P12 | -0.04 |
| Awili-01 | Pōhue | 56 | none | 157 | 1.000 | P05 | 0.09 |
| Awili-02 | Pōhue | 63 | none | 157 | 1.000 | P05 | 0.09 |
| Koloa-01 | Kamehame | 19 | none | 110 | 1.000 | P08 | 0.27 |
| Koloa-02 | Kamehame | 25 | none | 110 | 1.000 | P08 | 0.27 |

### Marker development

Because the power of genetic relatedness improves more with increased marker polymorphism and heterozygosity than the total number of markers (Sefc and Koblmüller 2009), this study sought to develop a panel of 100-300 polyallelic microhaplotype loci (see Gattepaille and Jakobsson 2012; McKinney et al. 2017; Baetscher et al. 2018) to capture genetic variation specific to Hawaiian hawksbill turtles. Candidate loci for a PCR-amplicon array were chosen from double-digested restriction site associated DNA sequences (ddRADseq; Peterson, Weber, Kay, Fisher, and Hoeskstra, 2012) from 18 Hawaiian hawksbills. Individual DNA concentrations were normalized to 500ng and samples were digested with two independent sets of two enzymes: set one (library 1) included EcoR1 and Sph1, and set two (library 2) included MluC1 and Sph1. Genomic fragments between 400-600bps were excised from a 2% Agarose gel. All other details of the library preparations were performed as in Peterson et. al (2012). Libraries were sequenced on an Illumina MiSeq sequencing platform using a paired-end approach and a 600 cycle sequencing kit. The pipeline STACKS v1.34 (Catchen et al. 2013) was used to demultiplex raw reads and identify allelic variants, requiring a minimum stack depth of 4(m), a distance of 4(M) allowed between stacks, and a distance of 4(n) allowed between catalog loci. A total of 63,610 unique genomic segments were found, of which 2,439 held at least two SNPs for at least 5 individuals. Loci were also screened for the density of polymorphisms at 100-300 bp intervals. Three hundred and seventeen genomic segments were randomly selected from the remaining Stacks loci for amplicon design, and PCR primers were designed for 259 of the candidate loci using FastPCR software (Kalendar et. al, 2017). Following 4 rounds of panel optimization, and the removal of paralogous sequences, 229 loci remained in the final panel.

### DNA extraction, PCR amplification, and DNA sequencing

Genomic DNA was extracted from tissue using a sodium chloride extraction (modified from Miller et al. 1988). Extracted DNA concentrations were normalized to 10 ng/ul. A total of 1,242 individuals were included for PCR amplification and sequencing, plus 180 replicates to assess genotyping consistency, and one negative control with no template DNA for every 96-well plate.

The Genotyping-in-Thousands by Sequencing protocol (GT-seq) of Campbell et al. (2015) was used to generate sequence data for genotype calling. An initial multiplex PCR containing locus-specific primers with Illumina (Illumina, Inc., San Diego, CA) priming sites for 229 amplicons was performed, followed by a second PCR to add Illumina adapters with indexes. Amplicon DNA concentrations were normalized across samples following PCR 2, using SequalPrep™ Normalization Plate Kits (Thermo Fisher Scientific), and pooled for a purification step performed using Agencourt ® AMPure ® XP magnetic beads to size select PCR products for sequencing. Each purified pool was quantified by Qubit Fluorometer (Thermo Fisher Scientific), and then by qPCR with the Illumina Library Quantification Kit (Kapa Biosystems). Pools were normalized to 4nM and pooled together. The library was sequenced on an Illumina Nextseq 500 using a single-end approach and a 150-cycle sequencing kit. All other details of the thermal cycling and library preparation are as in Campbell et al. (2015).

### Data processing, SNP genotyping, and microhaplotying

Adapter sequences, and base-pairs with a Phred quality score < 15, were trimmed from fastq reads using the program FASTP v. 0.23.1 (Chen et al. 2018). Sequences smaller than 90 bp after trimming were also excluded. Trimmed reads were then aligned to a fasta reference using the program BOWTIE2 v. 2.3.4.1 (Langmean and Salzburg 2012). Alignments were then sorted and indexed using SAMBAMBA v. 0.7.1 (Tarasov et al. 2015) while requiring a mapping quality score of ≥ 20. Sample alignments with fewer than 10,000 mapped reads were determined to have suboptimal amplicon sequencing depth and were excluded from further processing.

To reduce software biases introduced during genotyping (see Huwang et al. 2015; Ni et al. 2015; Sandmann et al. 2017; Ros-Freixedes et al. 2018) and improve genotyping accuracy (Jasper et al. 2022), variant calling was performed independently using three programs: BCFTOOLS v. 1.9 (Danecek et al. 2021), FREEBAYES v. 1.3 (Garrison and Marth 2012), and GATK-HC v. 3.8 (McKenna et al. 2010). Sensitivity was maximized for all three callers by accepting low alternate allele fractions (FREEBAYES), applying read fractions to individual rather than pooled samples (BCFTOOLS), and disabling pruning algorithms (GATK-HC). Called variants were reduced to their simplest components using the vcfallelicprimitives script from VCFLIB (Garrison et al. 2021). To minimize the tradeoff between genotyping sensitivity and accuracy, variants with low genotyping fidelity were identified using sequencing replicates and discarded if they had mismatched allele calls in > 7% of replicate sample pairs. In addition, variants were discarded if they weren’t identified by at least 2 out of the 3 variant callers, or had > 5.5% allelic mismatches among callers. The 7 and 5.5% mismatch thresholds were chosen after viewing histograms of allelic mismatches and making a qualitative determination about the level of genotyping noise (from unavoidable PCR and sequencing errors) that should be tolerated as normal (Supplementary Fig. 1). The custom R functions that performed the mismatch comparisons are available online (github.com/jh041/loc_gen_acc). SNPs and small indels that were robust to genotyping errors were then filtered for minor allele frequency and missing data using VCFTOOLS v. 0.1.16 (Danacek et al. 2011). Any variants with a minor allele frequency less than 0.01 were removed. Another experimental version of the data was also produced that required a minor allele frequency threshold of 0.05 and only allowed binary SNPs. Filtering for missing data followed an iterative approach, gradually decreasing missing data allowances from 80% to 30% for both loci and individuals. Variants from the three callers were filtered separately and not combined into a single data set until after microhaplotyping.

Microhaplotyping was performed using the R package MICROHAPLOT v. 1.0.1 (https://github.com/ngthomas/microhaplot) that uses both genotype calls and mapped reads to produce short phased haplotypes of all genetic variation found on each PCR amplicon. Additional filtering parameters were applied to microhaplotypes using MICROHAPLOT’s R Shiny app., including a minimum total microhaplotype read depth of 12, and an initial minimum allelic ratio of 0.50 (the minor microhaplotype allele must have a depth at least one half that of the major allele). Afterwards, the allelic ratios were refined for each amplicon locus individually by examining the relative depths of alleles for homo and heterozygous microhaplotype calls. The acceptable allele ratio for a homozygous call was never more than 0.09. The acceptable allele ratio for a heterozygous call was never less than 0.20. A minimum fraction of 0.7 microhaplotypes with acceptable allelic ratios across all individuals was required for each amplicon. If any individual had more than two possible microhaplotypes for the same locus (possibly indicating DNA contamination) the locus or sample was either removed, or noisy low-frequency microhaplotypes were excluded. At this point, microhaplotype calls arising from the FREEBAYES, BCFTOOLS, and GATK_HC outputs were combined into a single data set. Microhaplotype loci were removed from the analysis if more than 7% of allele calls were mismatched with replicate samples. Linkage among amplicon microhaplotypes (LD) and departures from Hardy-Weinberg expectations (HWE) were assessed using the program GENEPOP (Rousset 2008) and GENODIVE v. 3.05 (Meirmans 2020) adjusting *p*-values for multiple hypothesis testing using the method of Benjamini and Hochburg (1995).

### Genetic diversity and relatedness analysis

Genetic diversity statistics for our samples, including heterozygosity and *G*-statistics, were calculated for the final dataset using GENODIVE. Pairwise relatedness coefficients (*r*) were computed for all turtles with the R package RELATED v. 1.0 (Pew et al. 2015) using sample allele frequencies as reference points for the calculation of allelic states. The analysis is sensitive to genetic stratifications and linkage disequilibrium (Oliehoek et al. 2006), therefore, if two loci were shown to be in linkage disequilibrium then one of them was removed before relatedness analysis, and a number of different sample subsets were experimentally used to explore the sensitivity of the analysis to changes in the sample reference. The relative performances of the different relatedness estimators available from RELATED were evaluated using the native simulation modules for this package, generating 100 simulated genotypes each of four relatedness classifications (Parent-offspring, full-sibling, half-sibling, and unrelated). The estimator with the best correlation between simulated and inferred coefficients across all relatedness classes was then used to compute *r* for the empirical data. Final analysis was run with 1,000 bootstrap replicates to generate 95% confidence intervals for each pairwise value, and used an error-rate parameter of 0.02 for each locus. Inbreeding was set to “allowed.”

Pairwise relatedness was also inferred using the R package CKMRsim v. 0.1 (Anderson 2022), which uses a pseudo-likelihood approach and Monte Carlo pedigree simulations to model relatedness based on data-specific allele frequencies. In theory, this approach is less sensitive than relatedness coefficients to structures in the sample allele frequencies, because these become incorporated into the model, but power is still compromised by linkage disequilibrium between loci (Baetshcer et al. 2018). The results of this analysis were validated with 10,000 simulated genotype pairs of each relatedness type, generated using CKMRsim’s model framework. Simulated data was used to determine expected type-I and type-II error rates and baseline ranges of log-likelihood ratios for the following relatedness tests: parent-offspring vs. unrelated, full-sibling vs. unrelated, half-sibling vs. unrelated, and full-sibling vs. half-sibling. This analysis was also run with an assumed 2% error rate per locus.

Lastly, relatedness analysis was performed in COLONY v. 2.0.6.6 (Jones and Wang 2010), which differs from the other relatedness analyses by inferring the full pedigree likelihood of all samples simultaneously, instead of relying on pairwise inferences of relatedness. This analysis also uses pedigree information to assess genotyping error rates for each locus and reconstruct the pedigrees of unknown parents, such as the unsampled male hawksbill sires in this study. The paternal genotypes imputed from COLONY2 analysis were incorporated into all previously mentioned analyses as additional samples. Inbreeding and polygamy was allowed in the analysis. COLONY2 also estimates the effective population size (*N*_e_) of the breeding population using the sibship assignment method (Wang 2009).

Due to small population size, a strong signal of background inbreeding was suspected for Hawaiian hawksbill turtles, which can confound the accuracy of standard pairwise relatedness metrics (Brustad and Egeland 2019; Vigeland 2020). Though some of the used relatedness calculations have methods to reduce inbreeding biases implicit in the data (e.g. COLONY2), whether these were sufficient for the target population was not known *a priori*. However, because there were known relationships in our data (*i*.*e*. observed nesting females and offspring, full-sibling nestmates) we relied on relatedness inferences between these individuals to determine if inbreeding was adversely impacting relatedness estimates.

### Spatial patterns of genetic variation

In addition to relatedness inferences, several methods available in the R package ADEGENET (Jombart et al. 2011) were used to assess spatial allelic patterns using multivariate statistics that do not assume loci are in linkage equilibrium, or in Hardy-Weinberg proportions, and which are not sensitive to signals of selection or inbreeding. First, the data from 2017 nesting females and reconstructed paternal genotypes were clustered according to a *K*-means clustering algorithm, using 20 principal components as predictors. Discriminant Analysis of Principal Components (DAPC: Jombart et al. 2010) was then used to give a multivariate ordination of genetic differentiation based on these clusters. Finally, whether there was any positive spatial autocorrelation of genotypes among nesting sites was assessed using a spatial principal components analysis (Jombart et al. 2008). All individuals were georeferenced with latitude and longitude coordinates corresponding to nesting beaches where their offspring were born, with a slight amount of jitter added to avoid replicate coordinates. Statistical support for the result was determined using a Monte Carlo procedure (the global *r*test included in the ADEGENET package) and 1,000 permutations. The *p*-value of this test indicates the proportion of permuted statistics that exceed or are equal to the maximum observed value.

## Results

### Data processing, and genetic diversity

The mean number of raw, unmapped fastq DNA sequence reads per turtle was 366,235. Out of 1,422 raw fastq files, 257 were discarded for having less than 10,000 mappable reads, leaving 1,165 for analysis. The mean number of mapped reads across 1,165 samples was 271,490. The mean sequencing depth of PCR amplicons in the multiplex ranged from 22 to 7,500, with most having depths between 300 and 500. Of the 1,165 mapped samples 140 were replicates used to assess the consistency of genotype calls. The final data included 1,002 offspring from 41 nests, 13 adult females from the 2017 breeding season (Table 1), and 10 control females from the 2018 breeding season.

The three variant calling software programs each returned different numbers of raw SNP and indel DNA polymorphisms (BCFTOOLS = 1,839, FREEBAYES = 971, and GATK = 766). A total of 2,201 variants were detected by all programs. After removing variants with poor genotyping consistency across replicate samples and between variant calling programs, and filtering for missing data, there were 281, 202, and 256 variants from each of the callers, respectively. Only one of the final variants was an indel. Binary SNPs made up 85-98% of the other variants with the rest being trinary or quaternary SNPs. The final mean numbers of variants per PCR amplicon were 1.66, 1.72, and 1.58, for each of the callers respectively (Supplementary table 1). Preliminary analyses using only binary SNPs and a minor allele frequency threshold of 0.05 returned results consistent with the final data set that included all variant types.

A total of 170 PCR amplicons from all three callers were used for microhaplotyping and 135 final microhaplotype loci passed all filtering parameters, including linkage disequilibrium. The mean number of microhaplotype alleles per locus was 2.61, and the maximum number of alleles was seven (Fig. 1a). The mean missing data across all markers was 0.5% (Fig. 1b). The mean genotyping consistency of the final marker panel across replicate samples was 97.7%, and only two microhaplotype loci needed to be removed for having less than 93% consistency. Genotyping error rates estimated by COLONY2 suggest that the mean allelic drop rate across all markers was 3.5% (95% c.i = 1.77% - 6%), and the mean rate of all other errors was 0.4% (Fig. 1c,d). Estimated allelic drop rates were associated with the amount of missing data in each locus.

**Figure 1:**
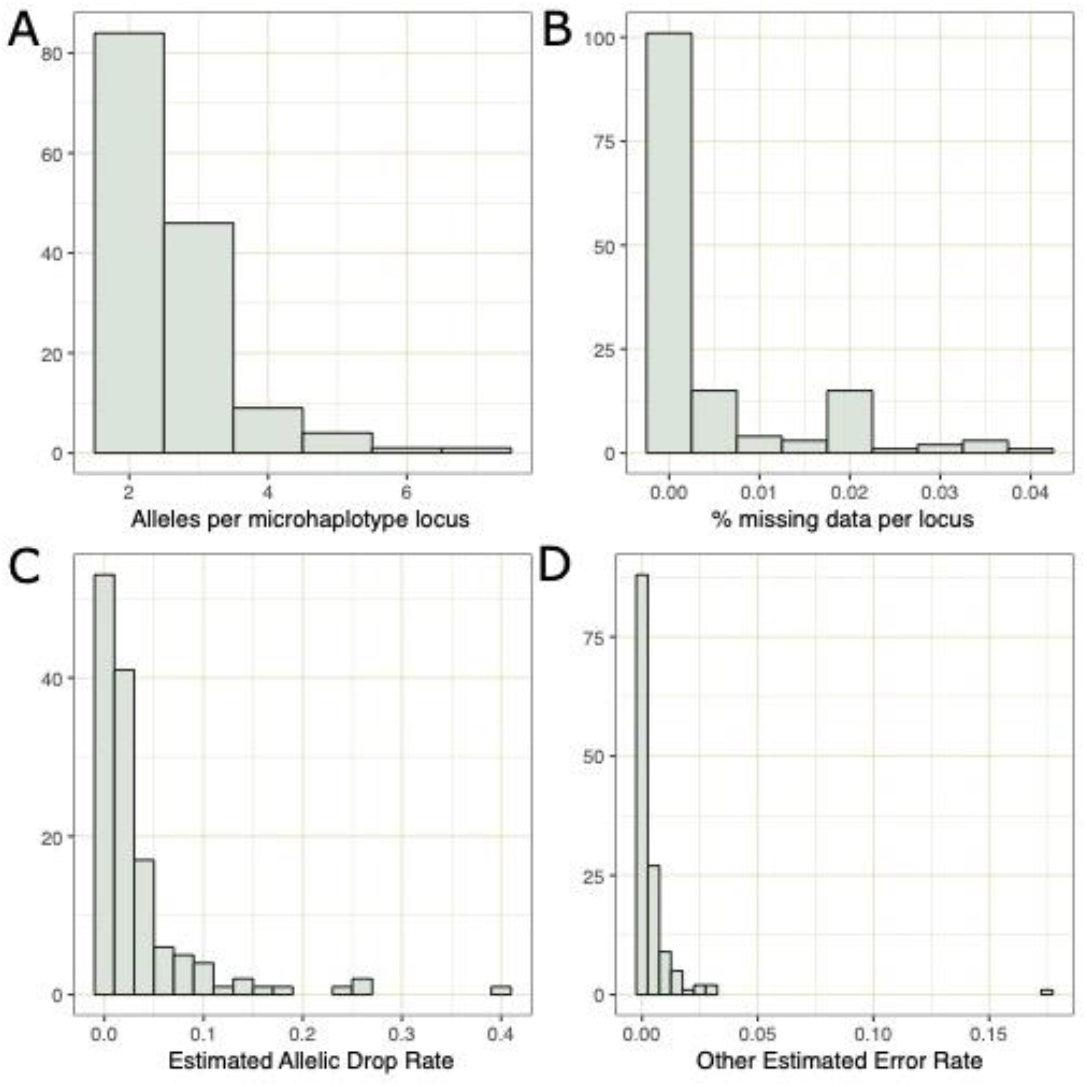
Descriptive statistics for 135 microhaplotye loci from 1,026 hawksbill turtles. **A)** Alleles per locus. **B)** The proportion of missing data per locus. **C)** Estimated allelic dropout rate per locus. **D)** Estimated rate of other errors per locus.

Patterns of genetic diversity differed between nesting complexes, especially across the southern point of the island (South Point) separating the ‘Āpua and Kamehame complexes in the east from the Pōhue complex in the west (Fig. 2a), with some loci having fixed alleles across this divide. Heterozygosity and allele frequencies were positively correlated for most loci between east and west nesting complexes, but not *G*_IS_ values (Fig. 3). The locus-specific *G*_IS_ values for both the east and west nesting complexes ranged between -0.7 and 0.7, with mean values falling below zero in both cases, and being significantly different from zero after 1,000 bootstrap replicates (Fig. 3c). In spite of many extreme *G*_IS_ values, departures from HWE were not statistically supported for any locus when both nesting complexes were analyzed jointly.

**Figure 2:**
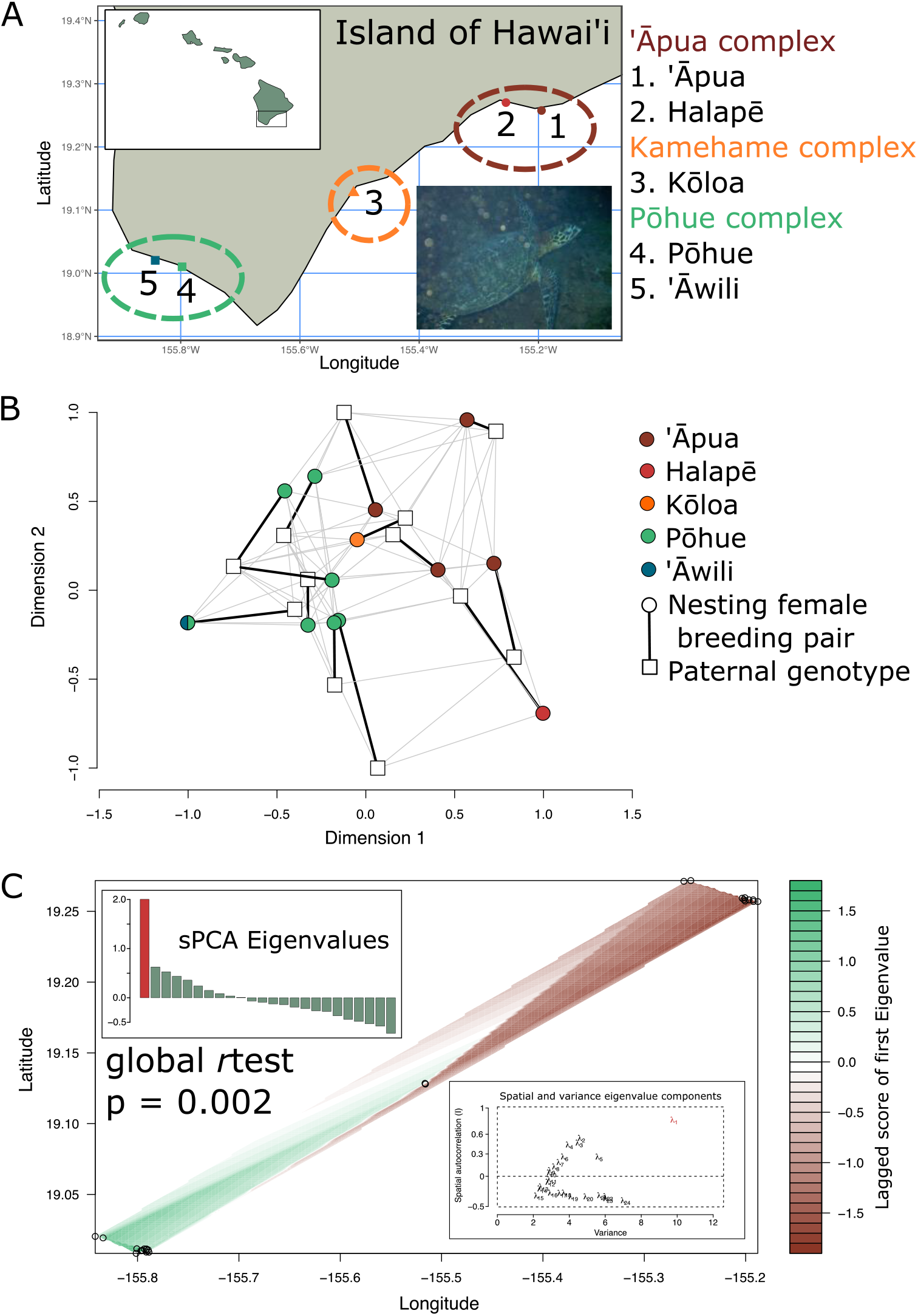
A) Map of the study area and the three nesting complexes. Photo credit: John B. Horne **B)** Relatedness network. Nodes represent nesting mothers and the inferred genotypes of their mates, reconstructed from hatchling microhaplotype loci. Edges are multi-dimensionally scaled pairwise relatedness values (1 - *r*). Edges smaller than *r* = 0 are omitted. **C)** Interpolated map of individual scores from spatial principal coordinates analysis. Inserts depict the relative contributions of each eigenvalue to the spatial autocorrelation of genetic variation.

**Figure 3:**
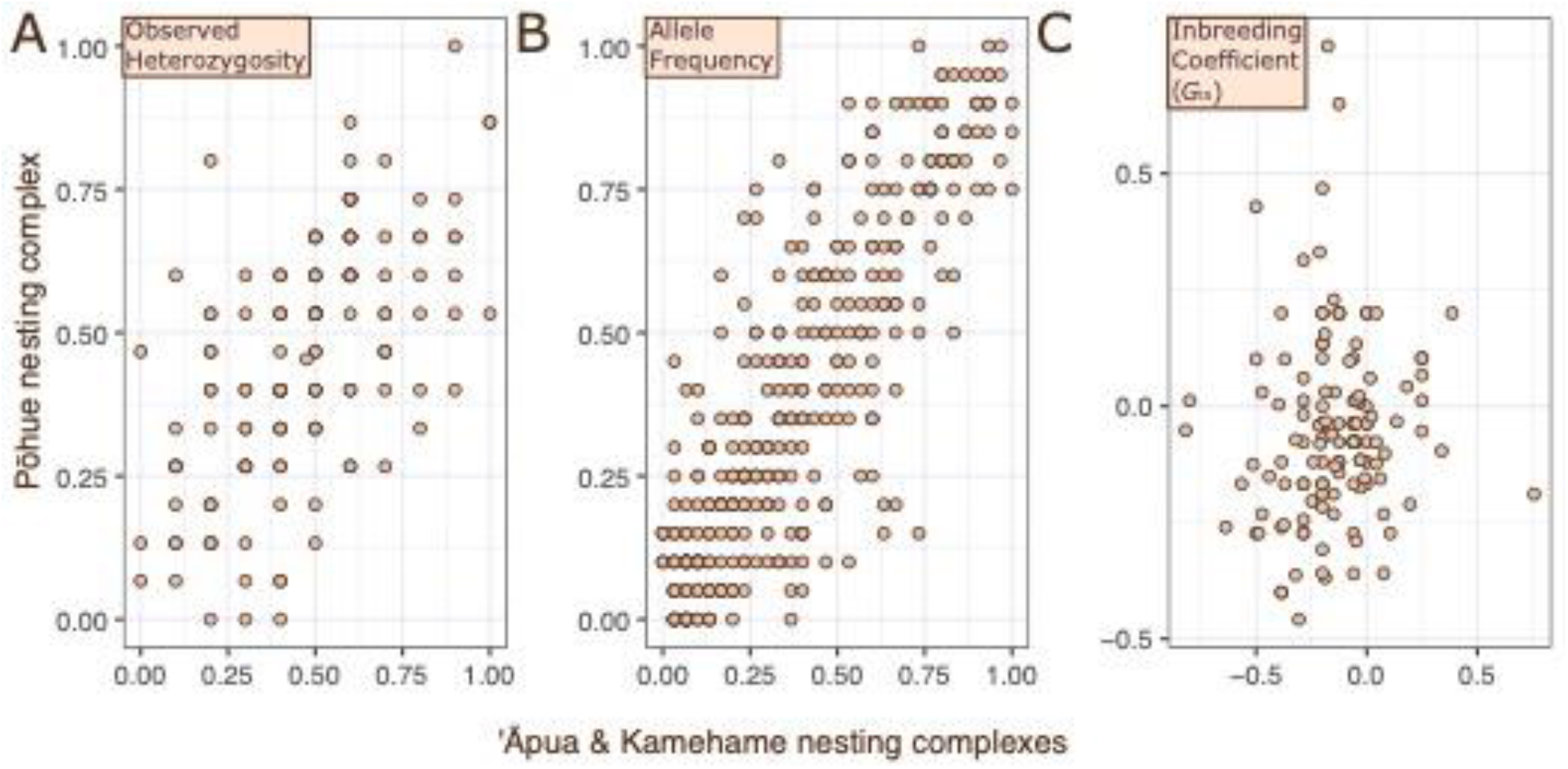
Population genetic comparisons of the Pohue nesting complex vs. the ‘Āpua and Kamehame nesting complexes: A) observed heterozygosity, B) allele frequency, C) the inbreeding coefficient *G*_IS_ of nesting females and inferred paternal male genotypes.

### Relatedness analysis

The best method-of-moments relatedness estimator for our data was that of Lynch and Ritland (1999), being over 94% correlated with the simulated levels of relatedness, and is hereafter referred to as *r*. The *r* distributions of known pairwise parent-offspring and full-sibling relationships from this study overlapped with simulated ranges but empirical *r* values tended to be lower than simulated data (Fig. 4). Changing the reference allele frequencies (by including or excluding offspring, 2018 nesters, or only using allele frequencies from the same nesting complex) did little to alleviate the skew. Nevertheless, whenever mean *r* was low the upper 95% confidence limit was helpful as a secondary measure for comparing pairwise relatedness to simulation expectations (Fig. 4).

**Figure 4:**
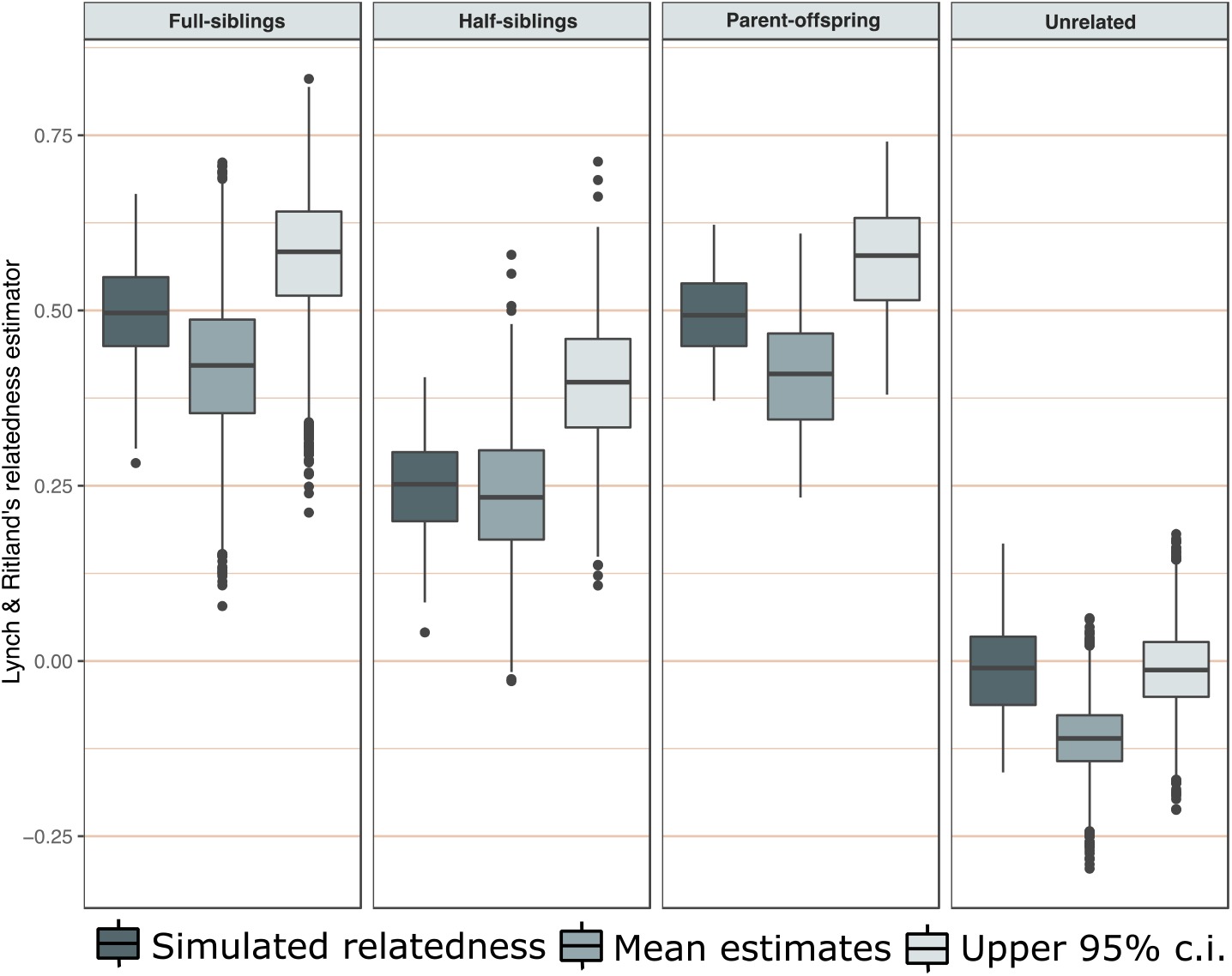
Boxplots showing the ranges of pairwise relatedness values (Lynch & Ritland 1999) for full-siblings, half-siblings, parent-offspring pairs, and unrelated individuals. Simulated ranges were calculated from 100 simulated pairs for each relationship type. Observed ranges of parent-offspring and full-sibling relatedness are from individuals with known relationships. Mean and 95% confidence intervals were produced after 1,000 bootstrap replicates.

Log-likelihood ratios for known parent-offspring and full-sibling relationships were approximated by simulated distributions (Fig. 5). In general, methods-of-moments and likelihood-based relatedness methods were in strong agreement with each other, and both were concordant with COLONY2 results. Every nesting female with a known nest was correctly identified by all three analyses as the mother of the offspring (Table 1). Likewise, offspring from nests with an unknown mother had clear parentage with a single candidate female in all but two cases (Table 1). The only exceptions were nests Pohue-24 and -25, where COLONY2 indicated mother-159 as the best mother for both with only 68.7% confidence, and the other methods were equally inconclusive. However, nest Pohue-25 is known to have been laid by mother-160, which was not sampled for this study but may be closely related to mother-159.

**Figure 5:**
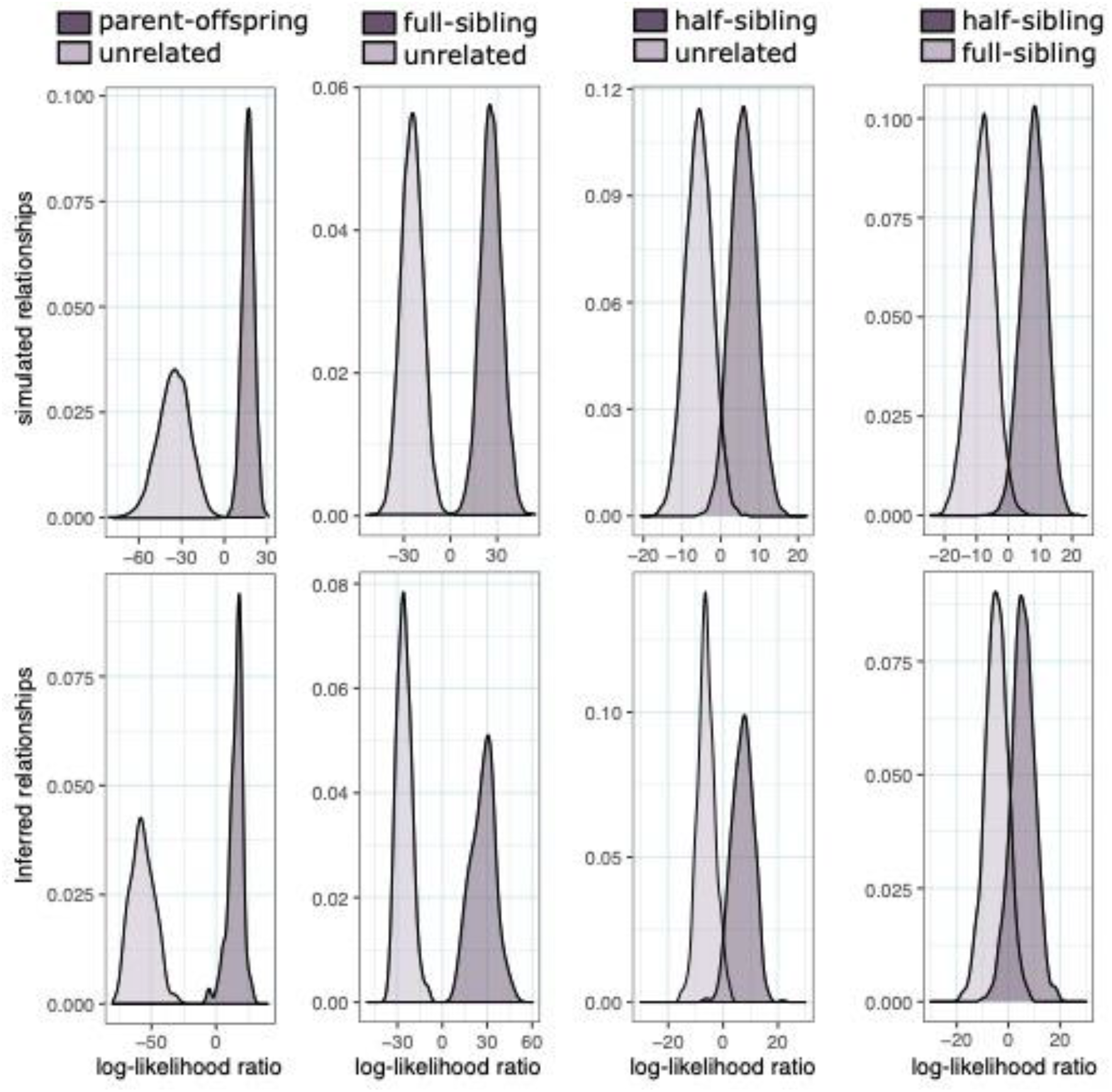
Kernel density plots of log-likelihood ratios for four different relationship comparisons. **Top row:** the expected log-likelihood ratio distributions for parent-offspring, full-sibling, and half-sibling pairs compared to unrelated individuals, from 10,000 simulated individuals for each category. **Bottom row:** observed log-likelihood ratios for each category. Distributions of parent-offspring and full-sibling ratios are from individuals with known relationships.

Given that nests Pohue-24 and -25 were likely laid by the same turtle, our data indicate 14 nesting females were responsible for the 41 sampled nests. Nesting females laid 1-5 nests with the mean being 2.9 (Table 1). Mother-157 was the only female to use multiple nesting sites within the same nesting complex, with two nests at ‘Āwili beach and one at Pōhue beach.

Paternal genotype reconstructions in COLONY2 indicate that there were 12 breeding males for 13 breeding females, with father P07 having mated with mother-119 and mother-176 (Table 1; Fig. 2b). Examination of pairwise relatedness between the offspring of these two nesting females corroborates this, indicating that they are half-siblings. Therefore, the operational breeding sex ratio for Hawaiian hawksbill turtles in 2017 appears to be near parity, with some female bias in the Pōhue nesting complex. There was no evidence of any half-sibling relationships among any of the nestmates, and no indication of polyandry during the 2017 breeding season. Mean estimates of N_e_ from COLONY2 were 8 and 16 for non-random and random mating scenarios respectively. The upper and lower 95% confidence bounds for N_e_ ranged from 4 to 34 across all estimates.

One unexpected result from the pairwise relatedness coefficients was that mean *r* for all the nesting females was noticeably less than the mean *r* values for the east and west nesting complexes individually. Closer inspection revealed that turtles breeding on opposite sides of South Point, both males and females, tended to be more related to each other than to turtles nesting across the point (Fig. 2b). Many breeding pairs also had high pairwise *r* values between them, especially from the ‘Āpua and Kamehame nesting complexes (Table 1). These data indicate that breeding individuals are mating non-randomly across the study area, and that inbreeding is not avoided when unrelated mates are available at other nearby breeding areas.

### Spatial patterns of genetic variation

The K-means clustering algorithm implemented in ADEGENET correctly split nesting females and inferred paternal genotypes between eastern and western nesting complexes, corroborating genetic diversity indices and relatedness analysis, and suggesting that these genetic differences are biologically meaningful. Genetic differences may also separate the eastern ‘Āpua and Kamehame nesting complexes, the latter of which was poorly sampled in the current study, but the available data indicate that the strongest genetic differentiation divides Hawai’i Island nesting turtles across South Point (Fig. 2). In terms of population structure, the *G*’_ST_ value between the eastern and western turtles was 0.036, p < 0.005; *G*’’_ST_ was 0.061, p < 0.005; and Jost’s *D* was 0.027, p < 0.005. When only the twelve most polymorphic microhaplotype loci, with four or more alleles, were assessed for population structure the indices were: *G*’_ST_ = 0.062, *G*’’_ST_ = 0.152, Jost’s *D* = 0.06, with all values being statistically supported at *p* < 0.005. In both sPCA and DAPC analysis a significant portion of the conserved genetic variance was spatially segregated across South Point (Fig. 3c). Thus, both male and female Hawaiian hawksbill turtles appear to exhibit strong natal philopatry and are sorting geographically according to nesting complex.

## Discussion

Though all sea turtles exhibit natal homing behaviors, genetic evidence shows variation among species, populations, and individuals in nesting site fidelity (Jensen et al. 2013). Female leatherback turtles (*Dermochelys coriacea*), for example, are not as strictly philopatric as other species (Thorson et al. 2012), and individuals display a range of nesting behaviors with some laying nests over 400 km apart in a single nesting season (Stewart et al. 2014b). Hawksbill females, in contrast, are among the more faithful of sea turtles to their natal nesting areas (Bowen et al. 2007; Monzón-Argüello et al. 2011; Gaos et al. 2016; Soanes et al. 2022), but many populations are genetically connected through male-mediated gene flow, particularly when rookeries are located along the same coastline (Phillips et al. 2014; Levasseur et al. 2019; see also Roden et al. in review). Compared to what has been reported for hawksbills in other parts of the world, the level of genetic differentiation observed among Hawaiian nesting complexes was high and unexpected.

Not only was there significant genetic population structure detected within the study area, but relatedness analysis revealed that both males and females appear to be mating assortatively by nesting complex (Fig. 2). The coastline distance between complexes (< 100 km) is trivial compared to the dispersal capabilities of an adult sea turtle, therefore, Hawaiian hawksbills appear to breed with other members of their same complex even when other mating opportunities are available. Contrasting signals of inbreeding and outbreeding among complexes at each locus (Fig. 3c) indicate that these results are not unique to the 2017 nesting season alone, because this pattern would require generations to form and would return mean *G*_IS_ values to zero after one generation of random mating. Population structure (*G*’_ST_) is likely exacerbated by small population size, but also indicates that there is limited longer-term genetic exchange. More data will be needed to make sense of the distinct east-west inbreeding patterns, but one explanation could be that Hawaiian hawksbill nesting colonies are currently trying to balance tradeoffs between inbreeding and outbreeding depression (Edmands 2007).

Inbreeding and outbreeding depression require an association between genetic diversity and reproductive fitness. In sea turtles, one measure of this fitness is likely the successful hatch rate of eggs. In the present study all sampled offspring were unsuccessful hatches, and the rate of successful hatches was assessed from empty egg shells in the nest chamber (data not shown). These data from the 13 females included in this study were not large enough for robust statistical analysis, but a study by Phillips et al. (2017) that was able to sample 95 nesting hawksbill females from the Seychelles, and their offspring, found evidence of both positive and negative correlations between multi-locus heterozygosity and hatching success, which suggests tension between inbreeding and outbreeding depression. Future studies of Hawaiian hawksbills, and other threatened populations of marine turtles, should pay closer attention to the relationship between genetic diversity and reproductive success, this being an aspect of their biology that is poorly understood but consequential for favorable conservation outcomes (Komoroske et al. 2017).

For breeding units as small as the nesting complexes of Hawaiian hawksbill turtles, even infrequent gene flow between them may be enough to prevent the loss of genetic diversity and stave off the worst effects of inbreeding (Whitlock et al. 2000; Whitley et al. 2015; Kardos et al. 2021; Sachdeva et al. 2022). This is because in a subdivided population random genetic drift will cause alleles to equilibrate differently in the different subunits, and the smallest subunits will experience the strongest genetic drift (Slatkin 1987). Gene flow between population subunits with different drift loads can yield substantial heterosis benefits, which are maximized when population sizes are small and migration between them is low (Whitlock et al. 2000). A loosely connected metapopulation of small breeding units would also theoretically be able to purge deleterious alleles more effectively, and potentially be more stable than a single randomly mating population with the same number of individuals (Sachdeva et al. 2022). Testing such a hypothesis for sea turtles would be difficult, but this could help explain why some populations appear perpetually small but steady over extended periods of time.

The mating system of Hawaiian hawksbills may also be important for genetic load management in this small population. This is because assortative mating within breeding groups can allow lineages to purge deleterious mutations more efficiently, in a similar fashion as subdivided populations (Muralidhar et al. 2022). In other parts of the world, hawksbill breeding sex ratios can be heavily female biased, presumably due to a limited supply of males in small populations (Gaos et al. 2018), but the Hawaiian rookeries have even fewer breeding individuals than elsewhere and a nearly 1:1 sex ratio, notwithstanding that females may outnumber males 4:1 overall in the archipelago (Brunson et al. 2022). Therefore, it cannot be ruled out that Hawaiian hawksbills are highly selective in their mate choices, even within nesting complexes (Fig. 2b; Table 1; see next section).

Because of the demographic discontinuity between nesting complexes there could in reality be multiple mating systems, and differences in breeding sex ratios, within the Hawaiian hawksbill metapopulation. At the outset of this research this degree of population complexity was not anticipated and more samples will be needed to elaborate on the patterns uncovered in this work. For example, more turtles from the Kamehame complex are required to determine genetic structure with nearby ‘Āpua (Fig. 2), and there are other nesting areas in the main Hawaiian islands that could be genetically and demographically unique. Most nesting sites not on the island of Hawaii are extremely low-density, having less than two nests laid annually, but one rookery on the island of Molokai has at least as many nesting females as any of the Hawaii Island nesting complexes (Gaos et al. 2021). Samples from multiple breeding seasons are also needed to better understand mating systems and female nesting behavior within and between complexes.

### Skew in estimates of pairwise relatedness

The genetic drift and mating system that determine the level of structure between two populations also determine the level of relatedness between two individuals, thus genetic population structure and genetic relatedness are two different aspects of the same variation (Weir and Goudet 2017). Just as measures of population structure can be seen as coancestry averages between the individuals in populations (Karhunen and Ovaskainen 2012), relatedness is a measure of coancestry between two individuals (Rousset 2002). The inextricable connection between genetic structure and genetic relatedness means that patterns in one are relevant to patterns in the other.

An overabundance of negative relatedness coefficients was observed for unrelated Hawaiian hawksbill turtles, and the *r* values of many related individuals were also depressed compared to distributions simulated under optimal conditions (Fig. 4). For the purposes of distinguishing related turtles from unrelated ones this skew is just noise, because all relatedness methods were able to correctly identify known relationships, regardless. But given the degree of inbreeding revealed by this study the skew deserves further exploration because negative pairwise *r* values are expected for samples that are outbred (have fewer loci that are identical-by-descent) relative to the reference allele frequencies (Wang 2014; 2017).

One source of skew in the data could be genetic structure in the reference allele frequencies, which can negatively bias relatedness estimates (Oliehoek et al. 2006; Wang 2017). Significant genetic structure was found between nesting complexes, but the skew in *r* values persisted even when the reference was generated exclusively from the same nesting complex. For the eastern nesting complexes, structure between Kamehame and ‘Āpua could be partly responsible, but the western Pōhue complex was also affected so other hidden structures could exist in the data. Family structures embedded within nesting complexes (or even nesting beaches) that create groups of highly related individuals could be the distortion. More data is required to determine if there could also be assortative mating within nesting complexes, as well as between them.

Relatedness coefficients can also be underestimated when the same samples for which relatedness is being estimated are used to estimate reference allele frequencies (Wang 2014, 2017). The strength of the bias is proportional to the number of samples included, on the order of 1/N where N is the number of samples. Therefore, when only the adult turtles in this study are used to generate the references we can expect the strength of the downward bias to be between 0.04 and 0.08. When including hatchlings, which theoretically have the same allele frequencies as their parents, the strength of the bias is between 0.01 and 0.001. Neither scenario fully explains the observed skew, though using hatchlings for the references might exacerbate negative biases due to family structures in the data.

Two more things that could be causing pairwise relatedness coefficients to be downwardly biased are selection and genetic admixture in the population founders (Wang 2014; Goudet et al. 2018). Balancing selection and purifying selection acting on different parts of the genome could be creating noise in pairwise relatedness estimates (Oliehoek et al. 2006), and the experimental loci do not necessarily need to be closely linked to the targets of selection because inbreeding is expected to reduce the rate of homologous recombination, creating large linkage blocks that are passed from generation to generation (Keller and Waller 2002; Charlesworth 2003). It is also plausible that hawksbills in the Hawaiian Islands are descended from multiple source populations, and if so then pedigrees would coalesce further back in time, creating a much deeper true reference relative to the present-day sample allele frequencies. How various population-level processes, such as inbreeding, are affecting pairwise relatedness estimates in hawksbill turtles would be clearer with more loci and a better understanding of the genomic architecture (see Kardos et al. 2015). Currently there is no genome sequence for *E. imbricata*, however, a Hawaiian hawksbill turtle was recently selected by the Vertebrate Genomes Project (vertebrategenomesproject.org) to represent this species as the reference genome, and this assembly will enable future research to explore the genomic complexity of this precariously small and non-randomly mating population.

## Acknowledgements

We recognize the Hawai’i Island Hawksbill Project, World Turtle Trust, and Hawai’i Volcanoes National Park for monitoring and skin sampling collection. This project received financial support from the National Oceanic and Atmospheric Administration Pacific Islands Regional Office under award numbers NA10, NA15, NA15NMF4540116, NA10NMF4540137, NA09NMF4540158, NA08NMF4540524, NA07NMF4540171, NA15NMF4540113, NA14NMF4540135, NA13NMF4540072, NA12NMF4540208. We also thank the United States Fish and Wildlife Service (USFWS) and the Hawai‘i State Department of Land and Natural Resources. We are indebted to Lauren Kurpita, Larry Katahira, Will Seitz, Joy Browning, Eldridge Naboa, Irene Kelly, T.Todd Jones, and the hundreds of dedicated volunteers that have participated in hawksbill monitoring at Hawai‘i Island over the years. Many thanks to Aliana Harmon, Erin LaCasella, Amy Lanci, Vicki Pease, and Gabriela Serra-Valente for their assistance with curation, processing, and laboratory analysis of samples accessioned into the Marine Mammal and Sea Turtle Research (MMSTR) Collection at the Southwest Fisheries Science Center’s La Jolla Laboratory in California. All hawksbill handling procedures were approved (SWPI_2013-05R) by the Institutional Animal Care and Use Committee in accordance with the requirements pertaining to animal subject protections within the Public Health Service Policy and United States Department of Agriculture Animal Welfare Regulations. Research was carried out under USFWS permits # TE8292509, TE739923, TE829250, TE739350, and TE72088A and State of Hawai’i Department of Land and Natural Resources Special Activity Permits.

## Tables and Figures

**Supplementary table 1a:** Data filtering: Raw genotyping results for 229 Hawksbill PCR amplicons from 1,032 Hawaiian Hawksbill turtles, plus 137 replicates.

| **Variant caller** | **FreeBayes** | **BCFtools** | **GATK** |
| --- | --- | --- | --- |
| Raw variants | 971 | 1839 | 766 |
| Raw binary SNPs | 696 | 1274 | 686 |
| Raw tri and quaternary SNPs | 235 | 268 | 19 |
| Raw indels | 40 | 297 | 61 |
| Raw PCR amplicons | 224 | 229 | 220 |
| Mean raw variants per PCR amplicon | 4.33 | 8.03 | 3.48 |

**Supplementary table 1b:** Data filtering: Variants with genotyping error rates of less than 7% (not including missing data). Error rates were calculated from mismatch tallies across 137 replicate sample pairs.

| **Variant caller** | **FreeBayes** | **BCFtools** | **GATK** |
| --- | --- | --- | --- |
| Mean raw allelic mismatches | 13.02 | 12.87 | 9.04 |
| Median raw allelic mismatches | 4 | 3 | 4 |
| Variants with < 7% genotyping error rate | 496 | 612 | 429 |
| PCR amplicons with < 7% genotyping error rate | 186 | 204 | 194 |
| Binary SNPs remaining | 306 | 441 | 402 |
| Tri and quaternary SNPs remaining | 161 | 93 | 15 |
| Indels remaining | 29 | 78 | 12 |
| Mean variants per PCR amplicon | 2.67 | 2.81 | 2.21 |

**Supplementary table 1c:** Data filtering: Low-error genotype variants from each caller that were also called by at least one other caller. Also, variants with less than 5.5% mismatched allele calls between callers across 137 replicate sample pairs.

| **Variant caller** | **FreeBayes** | **BCFtools** | **GATK** |
| --- | --- | --- | --- |
| Low-error variants identified by > 2 callers | 272 | 348 | 273 |
| Low-error PCR amplicons identified by > 2 callers | 138 | 174 | 161 |
| Variants called with < 5.5% errors between callers | 256 | 337 | 274 |
| PCR amplicons called with < 5.5% errors between callers | 128 | 163 | 151 |
| Binary SNPs remaining | 209 | 290 | 261 |
| Tri and quaternary SNPs remaining | 42 | 35 | 7 |
| Indels remaining | 5 | 12 | 6 |
| Mean variants per PCR amplicon | 2.00 | 1.97 | 1.82 |

**Supplementary table 1d:**
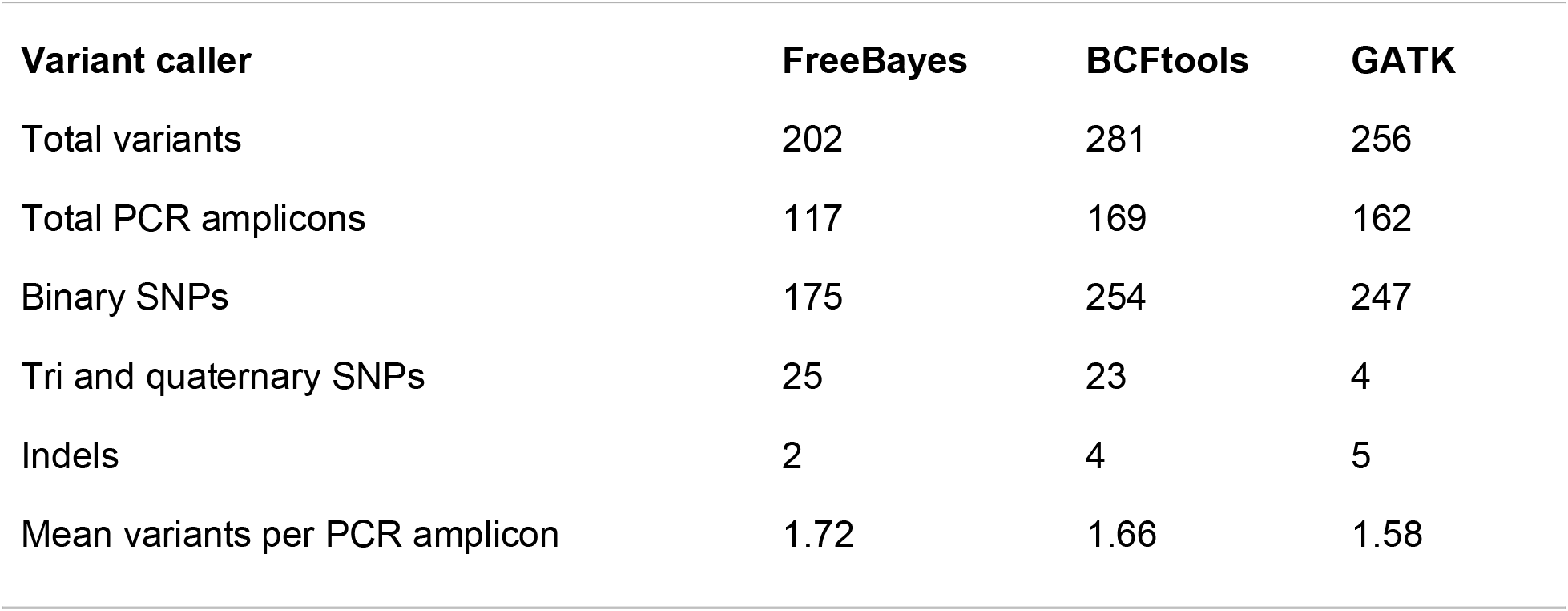
Data filtering: Variants remaining after filtering for > 30% missing data and a minor allele frequency < 0.01.

**Supplementary table 1e:** Data filtering: Microhaplotype calls before and after filtering. The filtration requirements were: a minimum sequence depth of 12, minimum heterozygote allelic ratio of 0.25, maximum homozygote allelic ratio of 0.09, and callable microhaplotypes in at least 70% of individuals.

| Variant caller | FreeBayes | BCFtools | GATK | Total |
| --- | --- | --- | --- | --- |
| Total microhaplotype loci | 117 | 169 | 162 | 170 |
| Filtered microhaplotype loci | 88 | 120 | 122 | 145 |
| Mean microhaplotype alleles per locus | 2.59 | 2.64 | 2.53 | 2.61 |
| Max microhaplotype alleles per locus | 6 | 7 | 6 | - |

**Supplementary figure 1:**
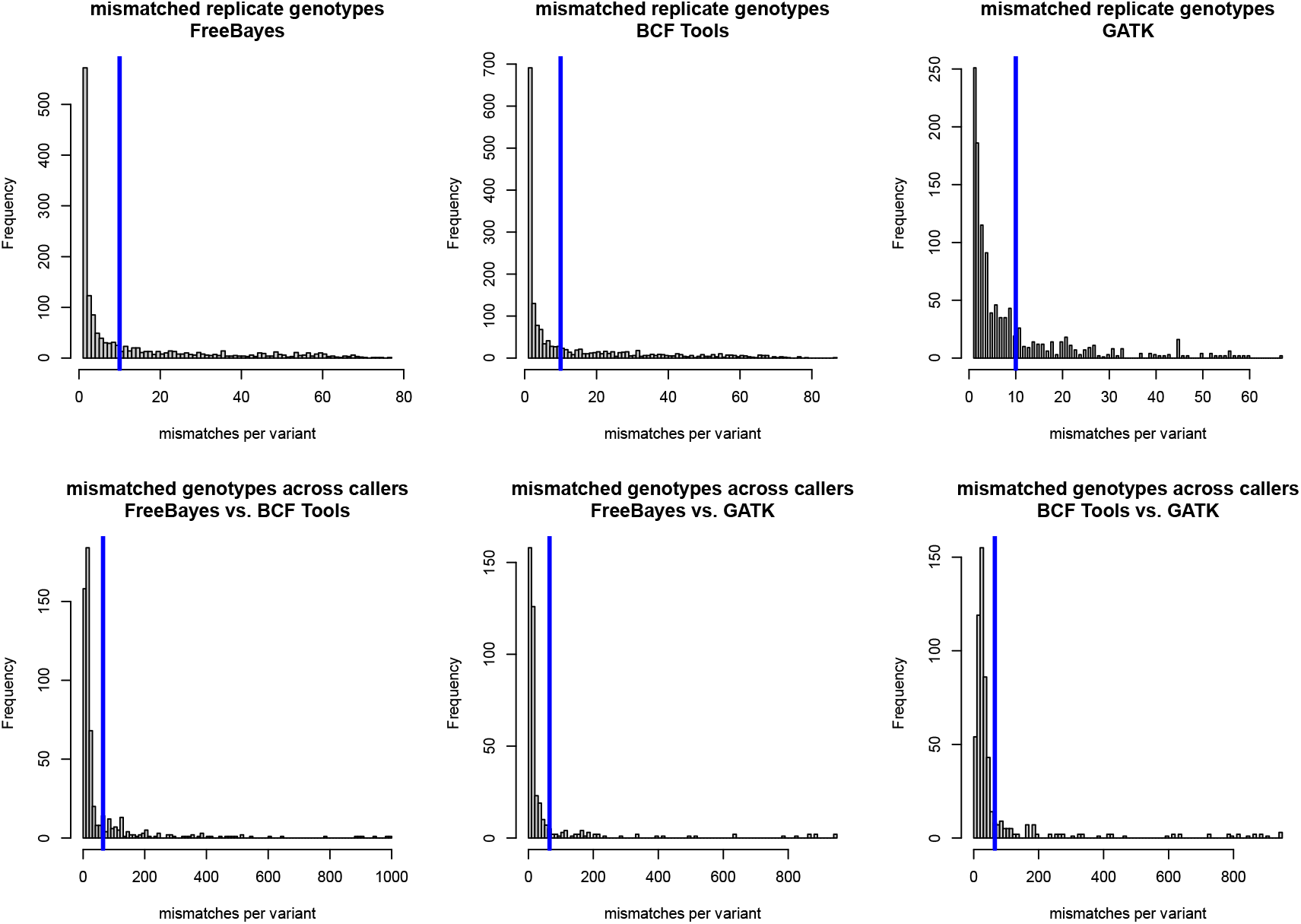
Histogram distributions of mismatches among genotype calls. Top row: mismatching single-locus genotype pairs across 137 replicate samples. Bottom row: mismatching single-locus genotypes across all variants called by two calling algorithms.Variants with mismatches exceeding the blue line thresholds (7% top, 5.5% bottom) were removed from the data sets.

## Notes

### Competing Interest Statement

The authors have declared no competing interest.

## References

Anderson, Eric C. 2022. “CKMRpop: Forward-in-Time Simulation and Tabulation of Pairwise Kin Relationships in Age-Structured Populations.” Molecular Ecology Resources 22 (3): 1190–99. https://doi.org/10.1111/1755-0998.13513.

Baetscher, Diana S., Anthony J. Clemento, Thomas C. Ng, Eric C. Anderson, and John C. Garza. 2018. “Microhaplotypes Provide Increased Power from Short-Read DNA Sequences for Relationship Inference.” Molecular Ecology Resources 18 (2): 296–305. https://doi.org/10.1111/1755-0998.12737.

Bell, I. P., J. J. Meager, T. Eguchi, K. A. Dobbs, J. D. Miller, and C. A. Madden Hof. 2020. “Twenty-Eight Years of Decline: Nesting Population Demographics and Trajectory of the North-East Queensland Endangered Hawksbill Turtle (Eretmochelys Imbricata).” Biological Conservation 241 (January): 108376. https://doi.org/10.1016/J.BIOCON.2019.108376.

Benjamini, Yoav, and Yosef Hochberg. 1995. “Controlling the False Discovery Rate: A Practical and Powerful Approach to Multiple Testing.” Journal of the Royal Statistical Society: Series B (Methodological) 57 (1): 289–300. https://doi.org/10.1111/j.2517-6161.1995.tb02031.x.

Bowen, B.W., W.S. Grant, Z. Hillis-Starr, J. Shaver, K.A. Bjorndal, A.B. Bolten, and A.L. Bass. 2007. “Mixed-Stock Analysis Reveals the Migrations of Juvenile Hawksbill Turtles (Eretmochelys Imbricata) in the Caribbean Sea.” Molecular Ecology 16: 49–60. https://doi.org/10.1111/j.1365-294X.2006.03096.x.

Brunson, Shandell, Alexander R. Gaos, Irene K. Kelly, Kyle S. Van Houtan, Yonat Swimmer, Stacy Hargrove, George H. Balazs, Thierry M. Work, and T. Todd Jones. 2022. “Three Decades of Stranding Data Reveal Insights into Endangered Hawksbill Sea Turtles in Hawai’I†.” Endangered Species Research 47 (February): 109–18. https://doi.org/10.3354/ESR01167.

Brustad, Hilde Kjelgaard, and Thore Egeland. 2019. “The Impact of Ignoring Inbreeding in Pairwise Kinship Evaluations.” Forensic Science International: Genetics Supplement Series 7 (1): 462–64. https://doi.org/10.1016/J.FSIGSS.2019.10.052.

Caballero, A., I. Bravo, and J. Wang. 2017. “Inbreeding Load and Purging: Implications for the Short-Term Survival and the Conservation Management of Small Populations.” Heredity 118 118: 177–85. https://doi.org/10.1038/hdy.2016.80.

Campbell, Nathan R., Stephanie A. Harmon, and Shawn R. Narum. 2015. “Genotyping-in-Thousands by Sequencing (GT-Seq): A Cost Effective SNP Genotyping Method Based on Custom Amplicon Sequencing.” Molecular Ecology Resources 15 (4): 855–67. https://doi.org/10.1111/1755-0998.12357.

Catchen, Julian, Paul A. Hohenlohe, Susan Bassham, Angel Amores, and William A. Cresko. 2013. “Stacks: An Analysis Tool Set for Population Genomics.” Molecular Ecology 22 (11): 3124–40. https://doi.org/10.1111/MEC.12354.

Chaloupka, Milani, Karen A. Bjorndal, George H. Balazs, Alan B. Bolten, Llewellyn M. Ehrhart, Colin J. Limpus, Hiroyuki Suganuma, Sebastian Troëng, and Manami Yamaguchi. 2008. “Encouraging Outlook for Recovery of a Once Severely Exploited Marine Megaherbivore.” Global Ecology and Biogeography 17 (2): 297–304. https://doi.org/10.1111/J.1466-8238.2007.00367.X.

Charlesworth, Deborah, and T. R. Meagher. 2003. “Effects of Inbreeding on the Genetic Diversity of Populations.” In Philosophical Transactions of the Royal Society B: Biological Sciences, 358:1051–70. The Royal Society. https://doi.org/10.1098/rstb.2003.1296.

Chen, Shifu, Yanqing Zhou, Yaru Chen, and Jia Gu. 2018. “Fastp: An Ultra-Fast All-in-One FASTQ Preprocessor.” Bioinformatics 34 (17): i884–90. https://doi.org/10.1093/BIOINFORMATICS/BTY560.

Danecek, Petr, Adam Auton, Goncalo Abecasis, Cornelis A. Albers, Eric Banks, Mark A. DePristo, Robert E. Handsaker, et al. 2011. “The Variant Call Format and VCFtools.” Bioinformatics 27 (15): 2156. https://doi.org/10.1093/BIOINFORMATICS/BTR330.

Danecek, Petr, James K Bonfield, Jennifer Liddle, John Marshall, Valeriu Ohan, Martin O Pollard, Andrew Whitwham, et al. 2021. “Twelve Years of SAMtools and BCFtools.” GigaScience 10 (2): 1–4. https://doi.org/10.1093/GIGASCIENCE/GIAB008.

Edmands, Suzanne. 2007. “Between a Rock and a Hard Place: Evaluating the Relative Risks of Inbreeding and Outbreeding for Conservation and Management.” Molecular Ecology. John Wiley & Sons, Ltd. https://doi.org/10.1111/j.1365-294X.2006.03148.x.

Frey, Amy, Peter H. Dutton, and George H. Balazs. 2013. “Insights on the Demography of Cryptic Nesting by Green Turtles (Chelonia Mydas) in the Main Hawaiian Islands from Genetic Relatedness Analysis.” Journal of Experimental Marine Biology and Ecology 442 (April): 80–87. https://doi.org/10.1016/J.JEMBE.2013.01.030.

Frey, Amy, Peter H. Dutton, Donna J. Shaver, Jennifer Shelby Walker, and Cynthia Rubio. 2014. “Kemp’s Ridley Lepidochelys Kempii Nesting Abundance in Texas, USA: A Novel Approach Using Genetics to Improve Population Census.” Endangered Species Research 23 (1): 63–71. https://doi.org/10.3354/ESR00565.

Gaos, Alexander R., Lauren Kurpita, Hannah Bernard, Luke Sundquist, Cheryl S. King, Joy H. Browning, Eldridge Naboa, et al. 2021. “Hawksbill Nesting in Hawai’i: 30-Year Dataset Reveals Recent Positive Trend for a Small, Yet Vital Population.” Frontiers in Marine Science 8 (December): 1719. https://doi.org/10.3389/FMARS.2021.770424/BIBTEX.

Gaos, Alexander R, Erin L Lacasella, Lauren Kurpita, · George Balazs, Stacy Hargrove, Cheryl King, Hannah Bernard, · T Todd Jones, · Peter, and H Dutton. 2020. “Hawaiian Hawksbills: A Distinct and Isolated Nesting Colony in the Central North Pacific Ocean Revealed by Mitochondrial DNA.” Conservation Genetics 21: 771–83. https://doi.org/10.1007/s10592-020-01287-1.

Gaos, Alexander R., Rebecca L. Lewison, Michael J. Liles, Ana Henriquez, Sofía Chavarría, Ingrid L. Yañez, Kelly Stewart, Amy Frey, T. Todd Jones, and Peter H. Dutton. 2018. “Prevalence of Polygyny in a Critically Endangered Marine Turtle Population.” Journal of Experimental Marine Biology and Ecology 506 (September): 91–99. https://doi.org/10.1016/j.jembe.2018.06.004.

Gaos, Alexander R, Rebecca L Lewison, Michael J Liles, Velkiss Gadea, Eduardo Altamirano, Ana V Henriquez, Perla Torres, et al. 2016. “Hawksbill Turtle Terra Incognita: Conservation Genetics of Eastern Pacific Rookeries.” Ecology and Evolution 6 (4): 1251–64. https://doi.org/10.1002/ece3.1897.

Garrison E, Marth G. Haplotype-based variant detection from short-read sequencing. arXiv preprint 1207.3907 [q-bio.GN] 2012

Garrison, Erik, Zev N. Kronenberg, Eric T. Dawson, Brent S. Pedersen, and Pjotr Prins. 2021. “Vcflib and Tools for Processing the VCF Variant Call Format.” BioRxiv, May, 2021.05.21.445151. https://doi.org/10.1101/2021.05.21.445151.

Gattepaille, Lucie M., and Mattias Jakobsson. 2012. “Combining Markers into Haplotypes Can Improve Population Structure Inference.” Genetics 190 (1): 159–74. https://doi.org/10.1534/GENETICS.111.131136.

Goudet, Jérôme, Tomas Kay, and Bruce S. Weir. 2018. “How to Estimate Kinship.” Molecular Ecology 27 (20): 4121–35. https://doi.org/10.1111/MEC.14833.

Hamann, M., M. H. Godfrey, J. A. Seminoff, K. Arthur, P. C.R. Barata, K. A. Bjorndal, A. B. Bolten, et al. 2010. “Global Research Priorities for Sea Turtles: Informing Management and Conservation in the 21st Century.” Endangered Species Research 11 (3): 245–69. https://doi.org/10.3354/esr00279.

Hamilton, Richard J., Tomas Bird, Collin Gereniu, John Pita, Peter C. Ramohia, Richard Walter, Clara Goerlich, and Colin Limpus. 2015. “Solomon Islands Largest Hawksbill Turtle Rookery Shows Signs of Recovery after 150 Years of Excessive Exploitation.” PLOS ONE 10 (4): e0121435. https://doi.org/10.1371/JOURNAL.PONE.0121435.

Hedrick, Philip W., and Aurora Garcia-Dorado. 2016. “Understanding Inbreeding Depression, Purging, and Genetic Rescue.” Trends in Ecology & Evolution 31 (12): 940–52. https://doi.org/10.1016/J.TREE.2016.09.005.

Hoban, Sean, Michael Bruford, Josephine D’Urban Jackson, Margarida Lopes-Fernandes, Myriam Heuertz, Paul A. Hohenlohe, Ivan Paz-Vinas, et al. 2020. “Genetic Diversity Targets and Indicators in the CBD Post-2020 Global Biodiversity Framework Must Be Improved.” Biological Conservation 248 (August): 108654. https://doi.org/10.1016/J.BIOCON.2020.108654.

Hoffmann, Ary A., Carla M. Sgrò, and Torsten N. Kristensen. 2017. “Revisiting Adaptive Potential, Population Size, and Conservation.” Trends in Ecology & Evolution 32 (7): 506–17. https://doi.org/10.1016/J.TREE.2017.03.012.

Hughes, A. Randall, and John J. Stachowicz. 2004. “Genetic Diversity Enhances the Resistance of a Seagrass Ecosystem to Disturbance.” Proceedings of the National Academy of Sciences of the United States of America 101 (24): 8998–9002. https://doi.org/10.1073/PNAS.0402642101.

Hwang, Sohyun, Eiru Kim, Insuk Lee, and Edward M. Marcotte. 2015. “Systematic Comparison of Variant Calling Pipelines Using Gold Standard Personal Exome Variants.” Scientific Reports 2015 5:1 5 (1): 1–8. https://doi.org/10.1038/srep17875.

Jamieson, Ian G., and Fred W. Allendorf. 2012. “How Does the 50/500 Rule Apply to MVPs?” Trends in Ecology & Evolution 27 (10): 578–84. https://doi.org/10.1016/J.TREE.2012.07.001.

Jasper, Russ J., Tegan Krista McDonald, Pooja Singh, Mengmeng Lu, Clément Rougeux, Brandon M. Lind, and Sam Yeaman. 2022. “Evaluating the Accuracy of Variant Calling Methods Using the Frequency of Parent-Offspring Genotype Mismatch.” Molecular Ecology Resources 00: 1–10. https://doi.org/10.1111/1755-0998.13628.

Jensen, Michael P., Camryn D. Allen, Tomoharu Eguchi, Ian P. Bell, Erin L. LaCasella, William A. Hilton, Christine A.M. Hof, and Peter H. Dutton. 2018. “Environmental Warming and Feminization of One of the Largest Sea Turtle Populations in the World.” Current Biology 28 (1): 154-159.e4. https://doi.org/10.1016/j.cub.2017.11.057.

Johnson, D W, J Freiwald, and G Bernardi. 2016. “Genetic Diversity Affects the Strength of Population Regulation in a Marine Fish.” Ecology 97 (3): 627–39.

Jombart, T., S. Devillard, A. B. Dufour, and D. Pontier. 2008. “Revealing Cryptic Spatial Patterns in Genetic Variability by a New Multivariate Method.” Heredity 2008 101:1 101 (1): 92–103. https://doi.org/10.1038/hdy.2008.34.

Jombart, Thibaut, Ismaïl Ahmed, and Alex Bateman. 2011. “Adegenet 1.3-1: New Tools for the Analysis of Genome-Wide SNP Data.” BIOINFORMATICS APPLICATIONS NOTE 27 (21): 3070–71. https://doi.org/10.1093/bioinformatics/btr521.

Jombart, Thibaut, Sébastien Devillard, and François Balloux. 2010. “Discriminant Analysis of Principal Components: A New Method for the Analysis of Genetically Structured Populations.” BMC Genetics 11 (1): 1–15. https://doi.org/10.1186/1471-2156-11-94/FIGURES/9.

Jones, Owen R., and Jinliang Wang. 2010. “COLONY: A Program for Parentage and Sibship Inference from Multilocus Genotype Data.” Molecular Ecology Resources 10 (3): 551–55. https://doi.org/10.1111/J.1755-0998.2009.02787.X.

Kalendar, Ruslan, Bekbolat Khassenov, Yerlan Ramankulov, Olga Samuilova, and Konstantin I. Ivanov. 2017. “FastPCR: An in Silico Tool for Fast Primer and Probe Design and Advanced Sequence Analysis.” Genomics 109 (3–4): 312–19. https://doi.org/10.1016/J.YGENO.2017.05.005.

Kardos, M., G. Luikart, and F. W. Allendorf. 2015. “Measuring Individual Inbreeding in the Age of Genomics: Marker-Based Measures Are Better than Pedigrees.” Heredity 115 (1): 63–72. https://doi.org/10.1038/hdy.2015.17.

Kardos, Marty, Ellie E. Armstrong, Sarah W. Fitzpatrick, Samantha Hauser, Philip W. Hedrick, Joshua M. Miller, David A. Tallmon, and W. Chris Funk. 2021. “The Crucial Role of Genome-Wide Genetic Variation in Conservation.” Proceedings of the National Academy of Sciences of the United States of America 118 (48). https://doi.org/10.1073/PNAS.2104642118.

Karhunen, Markku, and Otso Ovaskainen. 2012. “Estimating Population-Level Coancestry Coefficients by an Admixture F Model.” Genetics 192: 609–17. https://doi.org/10.1534/genetics.112.140871.

Keller, Lukas F., and Donald M. Waller. 2002. “Inbreeding Effects in Wild Populations.” Trends in Ecology & Evolution 17 (5): 230–41. https://doi.org/10.1016/S0169-5347(02)02489-8.

Komoroske, Lisa M., Michael P. Jensen, Kelly R. Stewart, Brian M. Shamblin, and Peter H. Dutton. 2017. “Advances in the Application of Genetics in Marine Turtle Biology and Conservation.” Frontiers in Marine Science 4 (JUN): 156. https://doi.org/10.3389/fmars.2017.00156.

LaCasella, Erin L., Michael P. Jensen, Christine A. Madden Hof, Ian P. Bell, Amy Frey, and Peter H. Dutton. 2021. “Mitochondrial DNA Profiling to Combat the Illegal Trade in Tortoiseshell Products.” Frontiers in Marine Science 7 (January): 1225. https://doi.org/10.3389/fmars.2020.595853.

Langmead, Ben, and Steven L Salzberg. 2012. “Fast Gapped-Read Alignment with Bowtie 2.” Nature Methods 2012 9:4 9 (4): 357–59. https://doi.org/10.1038/nmeth.1923.

Lasala, Jacob A., Colin R. Hughes, and Jeanette Wyneken. 2018. “Breeding Sex Ratio and Population Size of Loggerhead Turtles from Southwestern Florida.” PLoS ONE 13 (1): e0191615. https://doi.org/10.1371/journal.pone.0191615.

Levasseur, K E, S P Stapleton, and J M Quattro. 2021. “Precise Natal Homing and an Estimate of Age at Sexual Maturity in Hawksbill Turtles.” Animal Conservation 24 (3): 523–35. https://doi.org/10.1111/acv.12657.

Levasseur, Kathryn E., Seth P. Stapleton, Mykl Clovis Fuller, and Joseph M. Quattro. 2019. “Exceptionally High Natal Homing Precision in Hawksbill Sea Turtles to Insular Rookeries of the Caribbean.” Marine Ecology Progress Series 620 (June): 155–71. https://doi.org/10.3354/MEPS12957.

Levasseur, KE, SP Stapleton, MC Fuller, and JM Quattro. 2019. “Exceptionally High Natal Homing Precision in Hawksbill Sea Turtles to Insular Rookeries of the Caribbean.” Marine Ecology Progress Series 620 (June): 155–71. https://doi.org/10.3354/meps12957.

Luque, Gloria M., Chloé Vayssade, Benoît Facon, Thomas Guillemaud, Franck Courchamp, and Xavier Fauvergue. 2016. “The Genetic Allee Effect: A Unified Framework for the Genetics and Demography of Small Populations.” Ecosphere 7 (7): e01413. https://doi.org/10.1002/ecs2.1413.

Lynch, Michael, and Kermit Ritland. 1999. “Estimation of Pairwise Relatedness With Molecular Markers.” Genetics 152 (4): 1753–66. https://doi.org/10.1093/GENETICS/152.4.1753.

McKenna, Aaron, Matthew Hanna, Eric Banks, Andrey Sivachenko, Kristian Cibulskis, Andrew Kernytsky, Kiran Garimella, et al. 2010. “The Genome Analysis Toolkit: A MapReduce Framework for Analyzing next-Generation DNA Sequencing Data.” Genome Research 20 (9): 1297–1303. https://doi.org/10.1101/GR.107524.110.

McKinney, Garrett J., James E. Seeb, and Lisa W. Seeb. 2017. “Managing Mixed-Stock Fisheries: Genotyping Multi-SNP Haplotypes Increases Power for Genetic Stock Identification.” Canadian Journal of Fisheries and Aquatic Sciences 74 (4): 429–34. https://doi.org/10.1139/cjfas-2016-0443.

Meirmans, Patrick G. 2020. “Genodive Version 3.0: Easy-to-Use Software for the Analysis of Genetic Data of Diploids and Polyploids.” Molecular Ecology Resources 20 (4): 1126–31. https://doi.org/10.1111/1755-0998.13145.

Michael Reed, J, L Scott Mills, John B Dunning Jr, Eric S Menges, Kevin S M C Kelvey, Robert Frye, Steven R Beissinger, Marie-charlotte Anstett, and Philip Miller. 2002. “Emerging Issues in Population Viability Analysis.” Conservation Biology 16 (1).

Miller, S. A., D. D. Dykes, and H. F. Polesky. 1988. “A Simple Salting out Procedure for Extracting DNA from Human Nucleated Cells.” Nucleic Acids Research 16 (3): 1215. https://doi.org/10.1093/nar/16.3.1215.

Monzón-Argüello, C., N. S. Loureiro, C. Delgado, A. Marco, J. M. Lopes, M. G. Gomes, and F. A. Abreu-Grobois. 2011. “Príncipe Island Hawksbills: Genetic Isolation of an Eastern Atlantic Stock.” Journal of Experimental Marine Biology and Ecology 407 (2): 345–54. https://doi.org/10.1016/J.JEMBE.2011.07.017.

Mortimer, J. A., and M. Donnelly. 2008. Eretmochelys imbricata. IUCN Red List of Threatened Species 2011.1: https://www.iucnredlist.org.

Muralidhar, Pavitra, Graham Coop, and Carl Veller. 2022. “Assortative Mating Enhances Postzygotic Barriers to Gene Flow via Ancestry Bundling.” Proceedings of the National Academy of Science 119(30) https://doi.org/10.1073/pnas.

NMFS and USFWS. 2013. Hawksbill sea turtle (Eretmochelys imbricata) 5-year review: summary and evaluation. NOAA, National Marine Fisheries Service, Silver Spring, MD.

Ni, Guiyan, Tim M. Strom, Hubert Pausch, Christian Reimer, Rudolf Preisinger, Henner Simianer, and Malena Erbe. 2015. “Comparison among Three Variant Callers and Assessment of the Accuracy of Imputation from SNP Array Data to Whole-Genome Sequence Level in Chicken.” BMC Genomics 2015 16:1 16 (1): 1–12. https://doi.org/10.1186/S12864-015-2059-2.

O’Grady, Julian J., Barry W. Brook, David H. Reed, Jonathan D. Ballou, David W. Tonkyn, and Richard Frankham. 2006. “Realistic Levels of Inbreeding Depression Strongly Affect Extinction Risk in Wild Populations.” Biological Conservation 133 (1): 42–51. https://doi.org/10.1016/J.BIOCON.2006.05.016.

Oliehoek, Pieter A, Jack J Windig, Johan A.M. Van Arendonk, and Piter Bijma. 2006. “Estimating Relatedness between Individuals in General Populations with a Focus on Their Use in Conservation Programs.” Genetics 173 (1): 483–96. https://doi.org/10.1534/genetics.105.049940.

Peterson, Brant K., Jesse N. Weber, Emily H. Kay, Heidi S. Fisher, and Hopi E. Hoekstra. 2012. “Double Digest RADseq: An Inexpensive Method for De Novo SNP Discovery and Genotyping in Model and Non-Model Species.” PLOS ONE 7 (5): e37135. https://doi.org/10.1371/JOURNAL.PONE.0037135.

Pew, Jack, Paul H. Muir, Jinliang Wang, and Timothy R. Frasier. 2015. “Related: An R Package for Analysing Pairwise Relatedness from Codominant Molecular Markers.” Molecular Ecology Resources 15 (3): 557–61. https://doi.org/10.1111/1755-0998.12323.

Phillips, Karl P., Jeanne A. Mortimer, Kevin G. Jolliffe, Tove H. Jorgensen, and David S. Richardson. 2014. “Molecular Techniques Reveal Cryptic Life History and Demographic Processes of a Critically Endangered Marine Turtle.” Journal of Experimental Marine Biology and Ecology 455 (June): 29–37. https://doi.org/10.1016/j.jembe.2014.02.012.

Phillips, K. P., T. H. Jorgensen, K. G. Jolliffe, and D. S. Richardson. 2017. “Evidence of Opposing Fitness Effects of Parental Heterozygosity and Relatedness in a Critically Endangered Marine Turtle?” Journal of Evolutionary Biology 30 (11): 1953–65. https://doi.org/10.1111/jeb.13152.

Pinsky, Malin L., and Stephen R. Palumbi. 2014. “Meta-Analysis Reveals Lower Genetic Diversity in Overfished Populations.” Molecular Ecology 23: 29–39. https://doi.org/10.1111/mec.12509.

Pritchard, Adam M., Cheryl L. Sanchez, Nancy Bunbury, April J. Burt, Jock C. Currie, Naomi Doak, Frauke Fleischer-Dogley, et al. 2022. “Green Turtle Population Recovery at Aldabra Atoll Continues after 50 Yr of Protection.” Endangered Species Research 47 (March): 205–15. https://doi.org/10.3354/ESR01174.

Riskas, Kimberly A., Mariana M.P.B. Fuentes, and Mark Hamann. 2016. “Justifying the Need for Collaborative Management of Fisheries Bycatch: A Lesson from Marine Turtles in Australia.” Biological Conservation 196 (April): 40–47. https://doi.org/10.1016/j.biocon.2016.02.001.

Ros-Freixedes, Roger, Mara Battagin, Martin Johnsson, Gregor Gorjanc, Alan J. Mileham, Steve D. Rounsley, and John M. Hickey. 2018. “Impact of Index Hopping and Bias towards the Reference Allele on Accuracy of Genotype Calls from Low-Coverage Sequencing.” Genetics Selection Evolution 2018 50:1 50 (1): 1–14. https://doi.org/10.1186/S12711-018-0436-4.

Rousset, F. 2002. “Inbreeding and Relatedness Coefficients: What Do They Measure?” Heredity 2002 88:5 88 (5): 371–80. https://doi.org/10.1038/sj.hdy.6800065.

Rousset, François. 2008. “GENEPOP’007: A Complete Re-Implementation of the GENEPOP Software for Windows and Linux.” Molecular Ecology Resources 8 (1): 103–6. https://doi.org/10.1111/J.1471-8286.2007.01931.X.

Sachdeva, Himani, Oluwafunmilola Olusanya, and Nick Barton. 2022. “Genetic Load and Extinction in Peripheral Populations: The Roles of Migration, Drift and Demographic Stochasticity.” Philosophical Transactions of the Royal Society B: Biological Sciences 377 (1846). https://doi.org/10.1098/rstb.2021.0010.

Sandmann, Sarah, Aniek O. De Graaf, Mohsen Karimi, Bert A. Van Der Reijden, Eva Hellström-Lindberg, Joop H. Jansen, and Martin Dugas. 2017. “Evaluating Variant Calling Tools for Non-Matched Next-Generation Sequencing Data.” Scientific Reports 2017 7:1 7 (1): 1–12. https://doi.org/10.1038/srep43169.

Sefc, Kristina M., and Stephan Koblmüller. 2009. “Assessing Parent Numbers from Offspring Genotypes: The Importance of Marker Polymorphism.” Journal of Heredity 100 (2): 197–205. https://doi.org/10.1093/JHERED/ESN095.

Slatkin, Montgomery. 1987. “Gene Flow and the Geographic Structure of Natural Populations.” Science 236 (4803): 787–92. https://doi.org/10.1126/science.3576198.

Soanes, L.M., J. Johnson, K. Eckert, K. Gumbs, L.G. Halsey, G. Hughes, K. Levasseur, et al. 2022. “Saving the Sea Turtles of Anguilla: Combining Scientific Data with Community Perspectives to Inform Policy Decisions.” Biological Conservation 268 (April): 109493. https://doi.org/10.1016/J.BIOCON.2022.109493.

Spielman, Derek, Barry W Brook, and Richard Frankham. 2004. “Most Species Are Not Driven to Extinction before Genetic Factors Impact Them.” Proceedings of the National Academy of Science 101 (42): 15261–64. https://www.pnas.org.

Stewart, Kelly R., and Peter H. Dutton. 2014. “Breeding Sex Ratios in Adult Leatherback Turtles (Dermochelys Coriacea) May Compensate for Female-Biased Hatchling Sex Ratios.” PLoS ONE 9 (2): e88138. https://doi.org/10.1371/journal.pone.0088138.

Stewart, Kelly R., and Peter H. Dutton. 2011. “Paternal Genotype Reconstruction Reveals Multiple Paternity and Sex Ratios in a Breeding Population of Leatherback Turtles (Dermochelys Coriacea).” Conservation Genetics 12 (4): 1101–13. https://doi.org/10.1007/s10592-011-0212-2.

Stewart, Kelly R., Kelly J. Martin, Chris Johnson, Nicole Desjardin, Scott A. Eckert, and Larry B. Crowder. 2014. “Increased Nesting, Good Survival and Variable Site Fidelity for Leatherback Turtles in Florida, USA.” Biological Conservation 176 (August): 117–25. https://doi.org/10.1016/j.biocon.2014.05.008.

Tarasov, Artem, Albert J. Vilella, Edwin Cuppen, Isaac J. Nijman, and Pjotr Prins. 2015. “Sambamba: Fast Processing of NGS Alignment Formats.” Bioinformatics 31 (12): 2032–34. https://doi.org/10.1093/BIOINFORMATICS/BTV098.

Thorson, James T., André E. Punt, and Ronel Nel. 2012. “Evaluating Population Recovery for Sea Turtles under Nesting Beach Protection While Accounting for Nesting Behaviours and Changes in Availability.” Journal of Applied Ecology 49 (3): 601–10. https://doi.org/10.1111/j.1365-2664.2012.02143.x.

Valdivia, Abel, Shaye Wolf, and Kieran Suckling. 2019. “Marine Mammals and Sea Turtles Listed under the U.S. Endangered Species Act Are Recovering.” PLoS ONE 14 (1): e0210164. https://doi.org/10.1371/journal.pone.0210164.

Houtan, Kyle S. Van, Devon L. Francke, Sarah Alessi, T. Todd Jones, Summer L. Martin, Lauren Kurpita, Cheryl S. King, and Robin W. Baird. 2016. “The Developmental Biogeography of Hawksbill Sea Turtles in the North Pacific.” Ecology and Evolution 6 (8): 2378–89. https://doi.org/10.1002/ece3.2034.

Houtan, Kyle S. Van, John N. Kittinger, Amanda L. Lawrence, Chad Yoshinaga, V. Ray Born, and Adam Fox. 2012. “Hawksbill Sea Turtles in the Northwestern Hawaiian Islands.” Chelonian Conservation and Biology 11 (1): 117–21. https://doi.org/10.2744/CCB-0984.1.

Vigeland, Magnus Dehli. 2020. “Relatedness Coefficients in Pedigrees with Inbred Founders.” Journal of Mathematical Biology 81 (1): 185–207. https://doi.org/10.1007/S00285-020-01505-X/FIGURES/15.

Wallace, B.P., Andrew D. DiMatteo, Alan B. Bolten, M. Y. Chaloupka, Brian J. Hutchinson, F. Alberto Abreu-Grobois, Jeanne A. Mortimer, et al. 2011. “Global Conservation Priorities for Marine Turtles.” Edited by Steven J. Bograd. PLoS ONE 6 (9): 12–13. https://doi.org/10.1371/journal.pone.0024510.

Walsh, Matthew R., Stephan B. Munch, Susumu Chiba, and David O. Conover. 2006. “Maladaptive Changes in Multiple Traits Caused by Fishing: Impediments to Population Recovery.” Ecology Letters 9 (2): 142–48. https://doi.org/10.1111/J.1461-0248.2005.00858.X.

Wang, J. 2014. “Marker-Based Estimates of Relatedness and Inbreeding Coefficients: An Assessment of Current Methods.” Journal of Evolutionary Biology 27 (3): 518–30. https://doi.org/10.1111/JEB.12315.

Wang, J. 2017. “Estimating Pairwise Relatedness in a Small Sample of Individuals.” Heredity 2017 119:5 119 (5): 302–13. https://doi.org/10.1038/hdy.2017.52.

Wang, Jinliang. 2009. “A New Method for Estimating Effective Population Sizes from a Single Sample of Multilocus Genotypes.” Molecular Ecology 18 (10): 2148–64. https://doi.org/10.1111/J.1365-294X.2009.04175.X.

Weir, Bruce S, and Jérôme Goudet. 2017. “A Unified Characterization of Population Structure and Relatedness.” Genetics 206: 2085–2103. https://doi.org/10.1534/genetics.116.198424.

Whiteley, A. R., Fitzpatrick, S. W., Funk, W. C., & Tallmon, D. A. (2015). Genetic rescue to the rescue. Trends in Ecology & Evolution, 30(1), 42–49. https://doi.org/10.1016/J.TREE.2014.10.009

Whitlock, Michael C., Pär K. Ingvarsson, and Todd Hatfield. 2000. “Local Drift Load and the Heterosis of Interconnected Populations.” Heredity 84 (4): 452–57. https://doi.org/10.1046/j.1365-2540.2000.00693.x.

Worm, Boris, Edward B. Barbier, Nicola Beaumont, J. Emmett Duffy, Carl Folke, Benjamin S. Halpern, Jeremy B.C. Jackson, et al. 2006. “Impacts of Biodiversity Loss on Ocean Ecosystem Services.” Science 314 (5800): 787–90. https://doi.org/10.1126/SCIENCE.1132294/SUPPL_FILE/1132294.WORM.SOM.PDF.

Wyneken, Jeanette, Kenneth J. Lohmann, and John A. Musick. 2013. The Biology of Sea Turtles: Volume III. Edited by Jeanette Wyneken, Kenneth J. Lohmann, and John A. Musick. The Biology of Sea Turtles. Vol. 3. https://doi.org/10.1201/b13895.

